# Milliseconds matter: Biomechanical inverse dynamics analysis is highly sensitive to imperfect data synchronization

**DOI:** 10.1101/2024.06.24.600392

**Authors:** Koen K. Lemaire, Arthur D. Kuo

**Affiliations:** Faculty of Kinesiology, University of Calgary, Calgary, Alberta, Canada; Biomedical Engineering Graduate Program, University of Calgary, Calgary, Alberta, Canada; Department of Human Movement Sciences, Vrije Universiteit Amsterdam, Netherlands (current)

**Author notes:** Corresponding author: Koen Lemaire.

**Keywords:** Walking, motion capture, force plate, synchronization, residual force

## Abstract

Inverse dynamics analysis is the primary means of quantifying joint moments and powers from biomechanical data. The data are often combined from force plates, motion capture cameras, and perhaps body-worn inertial sensors, and must be temporally synchronized to avoid potentially large inverse dynamics errors. The principles behind the errors, and the sensitivities for movements such as human walking, have yet to be demonstrated. Here we quantify how inverse dynamics computations of joint moments, powers, and work are highly sensitive to temporal mis-synchronization. We do this with (1) a theoretical examination of inverse dynamics, supported by (2) a simulated multi-body jumping movement, and (3) experimental human walking data. The theoretical analysis shows that root-mean-square errors in joint powers increase linearly with temporal mis-synchronization, and increase more for faster movements. With the other analyses we quantify the specific amount. For example, for human walking at 1.25 m/s, an artificially induced 5 ms lag of force relative to motion resulted in a 29% root-mean-square error of the ankle joint moment. The corresponding error in ankle joint power was 58%, and five times as much for walking at 2.2 m/s. The residual force, a measure of internal inconsistency in the data, increased by almost 1% body weight for each millisecond of mis-synchronization for walking. These sensitivities are relevant because standard experimental equipment usually synchronizes only the recording of data, but not processing latencies internal to equipment, which can and do cause mis-synchronization on the order of ten milliseconds. Biomechanical data should be synchronized to within a few milliseconds, and with respect to physical stimulus, to yield accurate inverse dynamics analysis.

## Introduction

Inverse dynamics analysis (IDA) produces estimates of the moments and powers exerted about the body’s joints, usually through a combination of ground reaction forces and motion capture data. It is applied to a wide array of biomechanical tasks, including during locomotion and posture control (Cahouët et al., 2002; Cappozzo et al., 1975; Winter, 1983), and to generate reference data to drive forward dynamics simulations (Pizzolato et al., 2017; Veerkamp et al., 2021). Naturally, the accuracy of inverse dynamics analysis is limited by the accuracy of the force and motion input data (Hatze, 1978; Riemer et al., 2008; Silva and Ambrosio, 2004). But even if these data are otherwise accurate, imperfect synchronization of forces and motions will still introduce errors. It is therefore important to understand how, and to what degree, temporal mis-synchronization can degrade inverse dynamics analysis.

The importance of data synchronization for inverse dynamics analysis has long been recognized. Previous literature has included methods to perform synchronization (Miller, 2014; Rome, 1995; Yeadon and King, 1999), as well as observations of temporal mis-synchronization in standard laboratory equipment ranging from a few to a hundred milliseconds or more (Komisar et al., 2017). The effects of such latencies have been quantified for human running by Günther et al. (2005), by re-analyzing data with artificially introduced mis-synchronization. They found that 5 ms of mis-synchronization was sufficient to induce errors of 100% in the energy balance for each joint, and as much as 28 J at the hip. This high degree of sensitivity for running suggests that other tasks and movements may be similarly affected by mis-synchronization.

Our own biomechanics laboratory’s standard experimental equipment illustrates the synchronization issue. Like many laboratories, we synchronously record forces from a standard split-belt instrumented treadmill (Bertec Corp., Columbus Ohio) with frames of optical motion capture (PhaseSpace Inc., San Leandro, California). We tested this set-up by applying physical stimuli such as gentle taps and other motions, and found that the recorded forces lagged the corresponding motion capture by about 8 ms. The mis-synchronization seems attributable to internal processing latencies, for example numerical multiplications to convert raw sensor outputs into forces and moments, that are insensitive to force sampling rate (e.g., 960 Hz). In other words, the forces are outputted by our treadmill about 8 ms after they actually occurred. We believe this is not unique to our laboratory: Standard six-axis force treadmills and platforms may exhibit similar latencies that could affect biomechanical analyses.

There are several ways to determine the effect of mis-synchronization. The most straightforward way is to apply artificial mis-synchronization (after Günther et al., 2005) to experimental data. This can yield a relative but not absolute sensitivity, because perfect synchronization is not generally achievable in experimental data. Absolute error can, however, be evaluated with simulated movements where the degree of synchronization may be precisely controlled. Finally, it may also be helpful to examine the inverse dynamics analysis approach analytically, to theoretically relate the errors due to mis-synchronization to the properties of the input signals, using dynamics principles.

The purpose of the present report is to quantify the effect of mis-synchronization on inverse dynamics analysis in three parts. In Part I, we perform a simple theoretical analysis that quantifies the effects of mis-synchronization depending on properties of the movement. In Part II, we describe a forward simulation of a squat jump and standard experimental walking data, to which artificial mis-synchronization is applied. In Part III, we demonstrate the magnitude of the resulting errors. These theoretical, computational, and experimental demonstrations show how errors arise from mis-synchronization and are explainable by general principles.

### Part I. Theoretical analysis of synchronization error

A simple analysis reveals that the errors in forces and powers from inverse dynamics analysis should increase linearly with the temporal mis-synchronization. This is shown first with a sinusoidal motion, and second with a Fourier series representation of an arbitrary motion. For both, we add mis-synchronization to the dynamic equilibrium equations and show its effects on inverse dynamics analysis. For all analyses, we define the mis-synchronization delay Δ*t* as the temporal delay between force and motion measurements, defined as positive for forces lagging motion.

We first consider a point mass *M* undergoing one-dimensional sinusoidal movement at amplitude *A* and angular frequency *ω*. The position *x* of the mass is given by

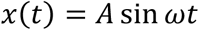

and the velocity *v(t)*, net force F*(t)*, and mechanical power *p(t)* are

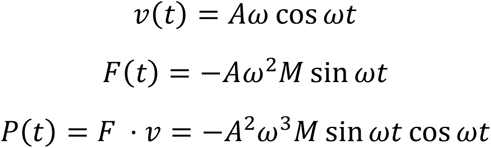

Suppose that *F(t)* and *x(t)* are recorded with perfect accuracy, except from a relative mis-synchronization Δ*t*. We wish to determine the effect of this mis-synchronization on the difference between the estimated and the veridical power. We denote the delayed recording *F*^d^*(t)* as

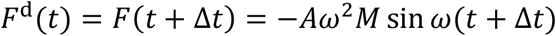

Applying a small angle approximation to *ω*Δ*t*, the difference between the delayed force and the original force is

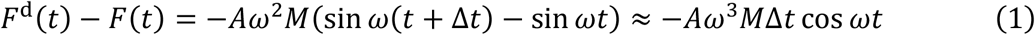

The force error therefore increases approximately linearly with mis-synchronization Δ*t*.

Based on this delayed force, the error between estimated power and veridical power becomes

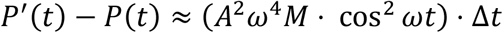

The root-mean-square error (RMSE) in mechanical power will thus become:

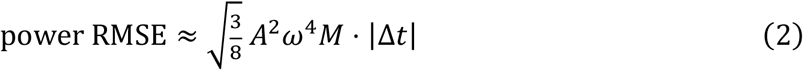

That is, the RMSE in the estimated mechanical power increases approximately linearly with mis-synchronization Δ*t*.

We next extend the same analysis to an arbitrary motion represented by a Fourier series. The motion is now

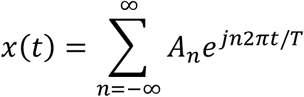

where *T* is either the period for periodic signals, or the duration of a non-periodic signal. (Signals that begin and end at different values may be adjusted by splicing together two copies, one reversed in time). The Fourier coefficients *A*_*n*_ are complex-valued, such that *x(t)* is real. The velocity *v(t)*, net force *F(t)*, and delayed force *F*^d^*(t)* are

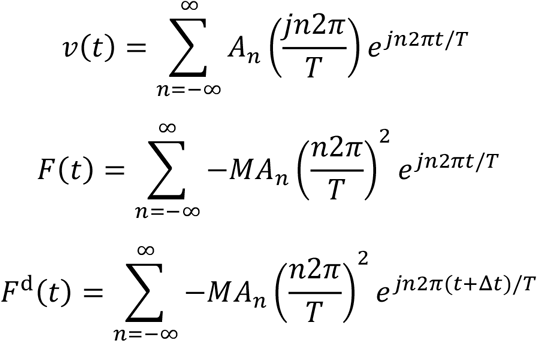

Following the same procedure of computing the mis-synchronized power and applying a small-angle approximation, the root-mean-square error in power has the form

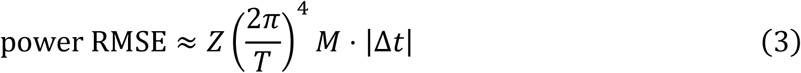

where *Z* is an amplitude-dependent quantity with units of squared amplitude. The number of terms in *Z* increases rapidly with the number of sinusoidal terms in *x(t)*. The fact that the overall expression is linear in Δ*t* may be understood from the property that sinusoids are mutually orthogonal, eliminating most of the cross-terms in the RMSE integral. Here, the small-angle approximation assumes small delay Δ*t* compared to the limited bandwidth of motion. High frequency motions can potentially violate this assumption, leading to even greater errors than reported here.

The above analysis shows that the root-mean-square power error increases approximately linearly with mis-synchronization Δ*t*. A body’s mechanical power increases linearly with mass, with the square of motion amplitude and with the cube of the motion frequency. The root-mean-square power error increases similarly, except with the fourth power of frequency. Thus, unsurprisingly, the error from mis-synchronization increases with mass and movement amplitude, and is particularly amplified for fast motions.

The errors due to mis-synchronization may be demonstrated with data in two ways. The first is in terms of the standard joint moments, powers, or work produced by inverse dynamics analysis. If the errors in resultant joint reaction forces increase approximately linearly with mis-synchronization delay, these errors should propagate into joint moment estimates, which depend linearly on the forces. The absolute errors may be quantified with respect to perfect synchronization in simulated data. For experimental data, only relative errors can be quantified, with respect to imperfect, nominal synchronization. Nevertheless, the theoretical analysis shows that errors should increase approximately linearly with mis-synchronization Δ*t*.

A second means to demonstrate mis-synchronization is in terms of residual forces and moments. Inverse dynamics analysis usually starts from ground reaction forces and proceeds upwards through the body segments, yielding residual forces and moments acting on the top-most body segment. These residuals should theoretically be zero, but in practice have non-zero values that are indicators of experimental error or imperfect modeling assumptions such as segment rigidity (van der Zee and Kuo, 2021). Just as with inter-segment forces, residual forces and moments will also be sensitive to Δ*t*. Any non-zero residual force is treated as error, and quantified as a function of Δ*t*.

### Part II. Simulation of jumping and Experimental data from walking

We describe two sets of example data used to demonstrate errors due to mis-synchronization. The first is a simulation of a squat jump, for which the ground truth synchronization is available. The second is experimental data from human walking. These represent relatively standard movements on which inverse dynamics analysis is typically applied. Both movements have been described previously in the literature, and are therefore only briefly summarized here.

### Forward simulation of squat jump

We used a previously published musculoskeletal model of maximum height squat jumping (van Soest and Bobbert, 1993) to generate human-like data. A simulation was done for the take-off phase of a squat jump, where there is considerable mechanical work performed. In contrast to the point-mass motions of the theoretical analysis, the simulation involves multibody dynamics. We considered the planar, skeletal part of the model, which consists of four linked, rigid segments representing the feet, shanks, thighs, and upper body (Fig. 1A). We used as inputs the initial model configuration and joint moment trajectories that maximize jump height (van Soest and Bobbert, 1993). The simulation includes the take-off phase only, starting initially standing still, and varying over time until lift-off (at time *t* = 0). The joint moments yielded a well-coordinated jump with the center of mass reaching a height of approximately 42 cm above standing. Mis-synchronization was applied by delaying the ground reaction force with respect to the nominal motion, again with positive mis-synchronization Δ*t* defined as force lagging motion data (Fig. 1 B/C). Inverse dynamics analysis was performed for mis-synchronization ranging from −15 ms to +15 ms. Results were non-dimensionalized using the model’s mass *M* (82 kg), leg length *L* (0.94 m) and the gravitational acceleration constant *g* (9.81 m/s^2^).

**Fig. 1.**
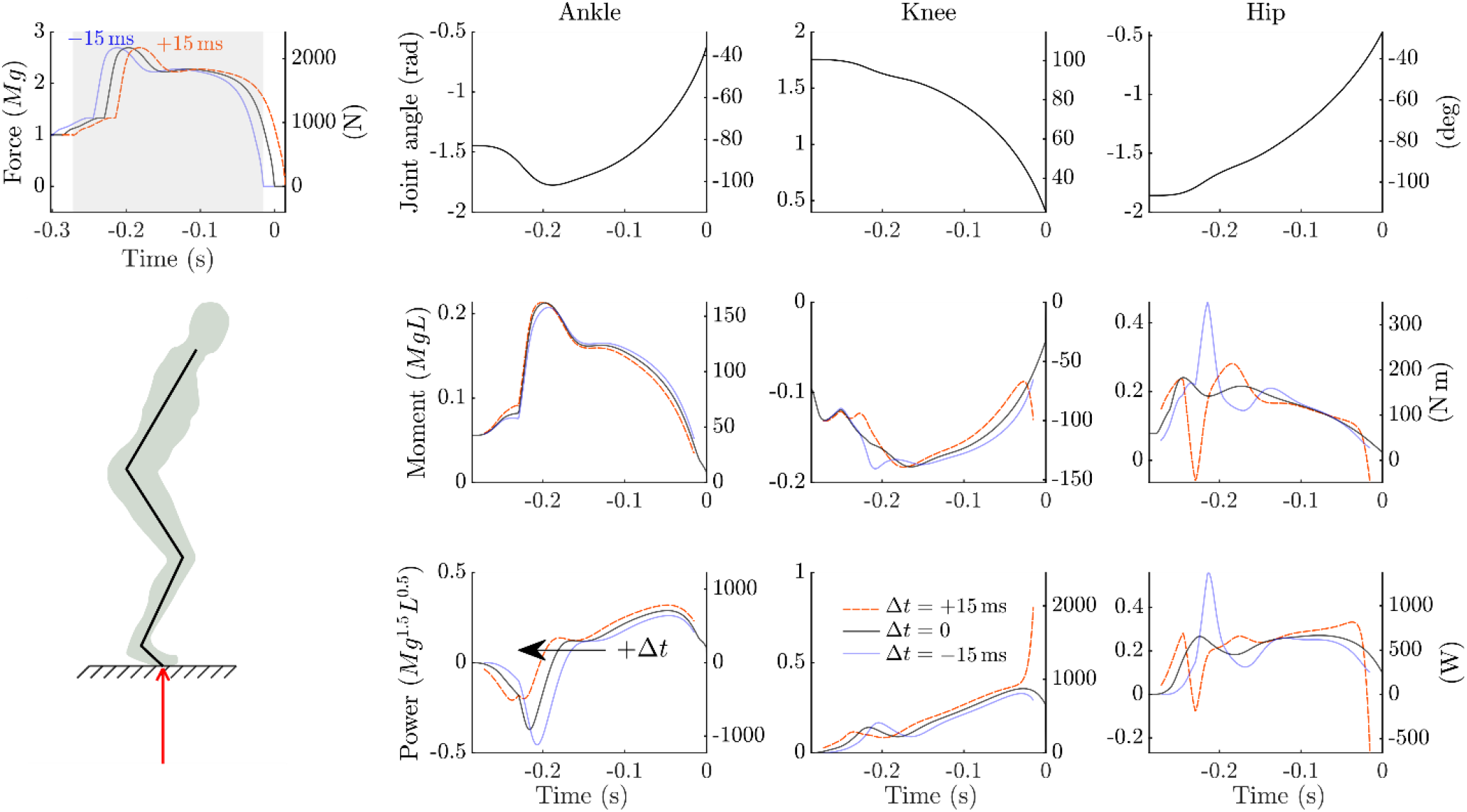
An optimal control simulation of jumping with mis-synchronization between forces and motions. The planar model (Bottom left) consisted of 4 segments representing the feet, shanks, thighs, and head-arms-trunk. The only external forces were the ground reaction force (red arrow) and gravity. The simulation with optimal joint moments (black solid lines) was performed for the take-off phase of the jump, yielding ground reaction force vs. time (Top left, only vertical component shown), as well as joint angle trajectories for ankle, knee, and hip (top row, right). Artificial delay was applied to the ground reaction forces, for mis-synchronization *Δt* of ±15 ms, defined as positive (red, dashed lines) for force lagging motion and negative (blue lines) for force leading motion. The shaded area indicates the interval for which the mis-synchronized data was analyzed, avoiding boundary effects beyond the time range of the simulation. Inverse dynamics analysis was performed on these data, yielding (rows below joint angles) joint moments and mechanical powers for the ankle, knee and hip joints vs. time. Results are shown for three values of *Δt* (±15 ms, 0 ms). Left-hand vertical axes are in normalized units and right-hand axes are in SI-units.

### Experimental data from human walking

We collected human walking data (Fig. 2) of one representative adult subject at a range of speeds (1.1 to 2.2 m/s) on an instrumented force treadmill, with standard motion capture recordings (c.f. Zelik and Kuo, 2010). Standard inverse dynamics analysis was performed to yield estimates of joint moment, power, and work. We again introduced artificial mis-synchronization of ground reaction forces and moments with respect to the kinematic data, and quantified the effect on the outcomes of inverse dynamics analysis. We used data from a 1.25 m/s walking speed as a nominal reference (see Fig. 2 for example data), and also examined errors as a function of walking speed, at a mis-synchronization of +5 ms. Data from only one subject was collected, because these data are intended to demonstrate errors due to mis-synchronization, and not to draw conclusions regarding human movement. We used body mass *M* (75 kg), leg length *L* (0.91 m), and gravitational acceleration *g* (9.81 m/s^2^) as normalizing units.

**Fig. 2.**
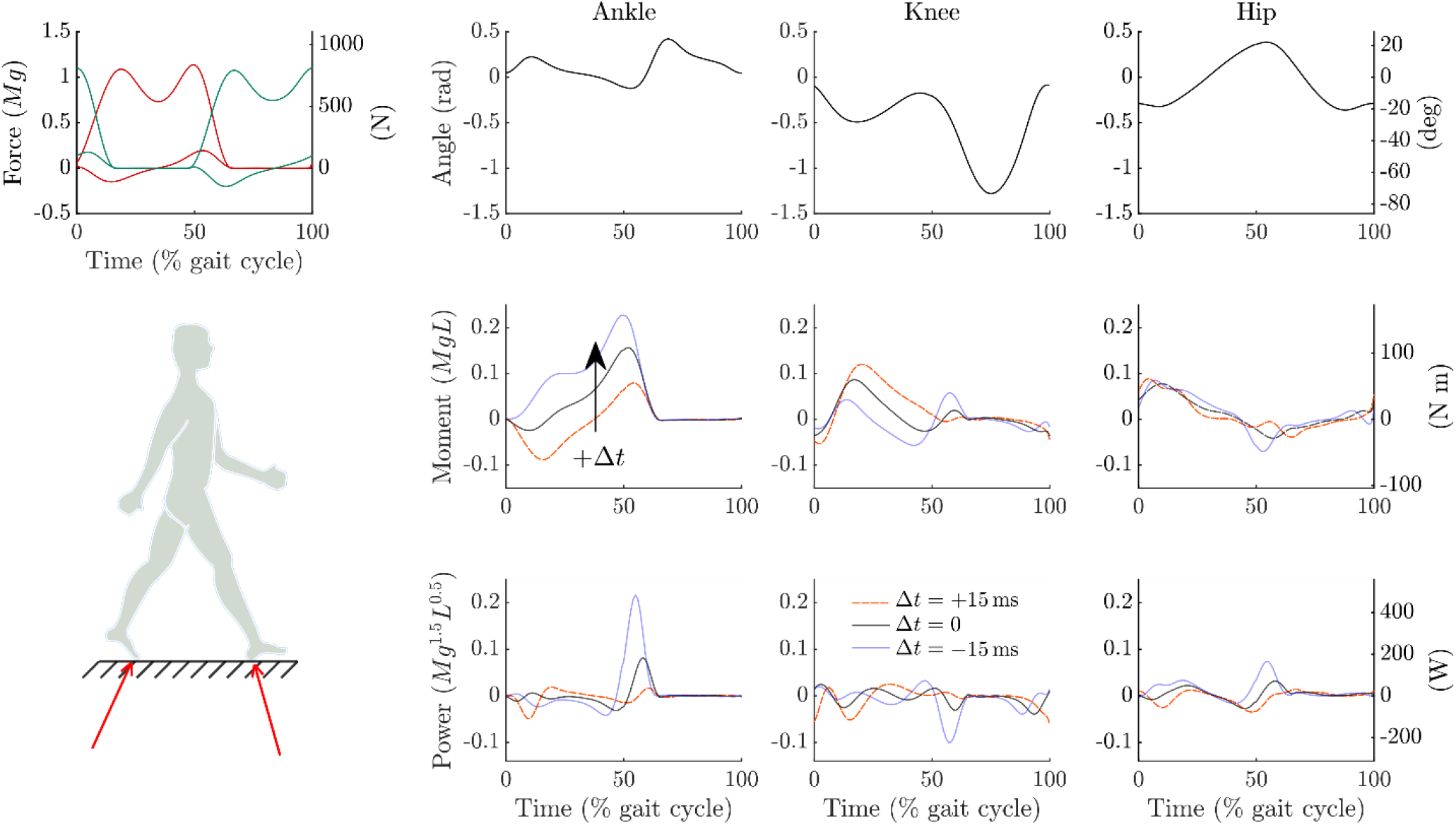
Experimental data for human walking with mis-synchronization between forces and motions. Data were collected from one representative adult subject walking at 1.25 m/s, including 3D ground reaction force vs. time (top left, vertical and fore-aft direction shown; left leg in red, right leg in green), as well as (top row, right) joint angle trajectories for ankle, knee, and hip (sagittal plane data shown), averaged over 42 consecutive strides. Artificial mis-synchronization *Δt* of ±15 ms was applied to the ground reaction forces, where positive (red, dashed) is defined as force lagging motion and negative (blue) as force leading motion. Inverse dynamics analysis was performed on these data, yielding (rows below joint angles) joint moments and mechanical powers for the ankle, knee and hip joints vs. time. Results are shown for three values of *Δt* (±15 ms, 0 ms). Left-hand vertical axes are normalized units and right-hand axes are SI-units; same axes are shared across all three joints.

The inverse dynamics analysis yielded joint angles, moments, and powers for the ankle, knee, and hip joints. We averaged data over the left and right sides of the body, with contralateral side shifted by 50% gait cycle, and averaged over 42 complete strides (Fig. 2). The resulting averaged lower limb kinematics and kinetics for the nominal case were qualitatively similar to previously published reports (Zelik and Kuo, 2010). We computed sensitivity of inverse dynamics analysis with respect to artificially induced mis-synchronization, treating as reference the original experimental data, unshifted in time. Although imperfect, this reference is sufficient to illustrate sensitivity to poor synchronization.

### Part III. Quantification of inverse dynamics errors due to mis-synchronization

We next quantify the effect of mis-synchronization on inverse dynamics analysis for both the squat jumping simulation and experimental human walking data. Based on theory (Part I), we expected an approximately linear increase in root-mean-square error for joint moments and powers, as a function of magnitude of delay Δ*t*. In the case of the squat jumping simulation, the mis-synchronization is relative to perfect ground truth of Δ*t* = 0. For experimental walking data, Δ*t* = 0 represents our best attempt at correct synchronization, found by shifting forces earlier in time by an experimentally tested latency of 8 ms for our equipment. A linear increase in root-mean-square error was expected with respect to that reference. For both data sets, all joint level values are reported for one single leg, represented by the average of the two legs.

The error introduced by the delay was substantial, in terms of both joint moments and powers (Fig. 3, top and middle rows). The root-mean-square errors (RMSE, with respect to ground truth) in both joint moments and powers increased approximately linearly with delay magnitude, for both positive and negative delays and for both jumping and walking (Fig. 3, left and right columns respectively).

**Fig. 3.**
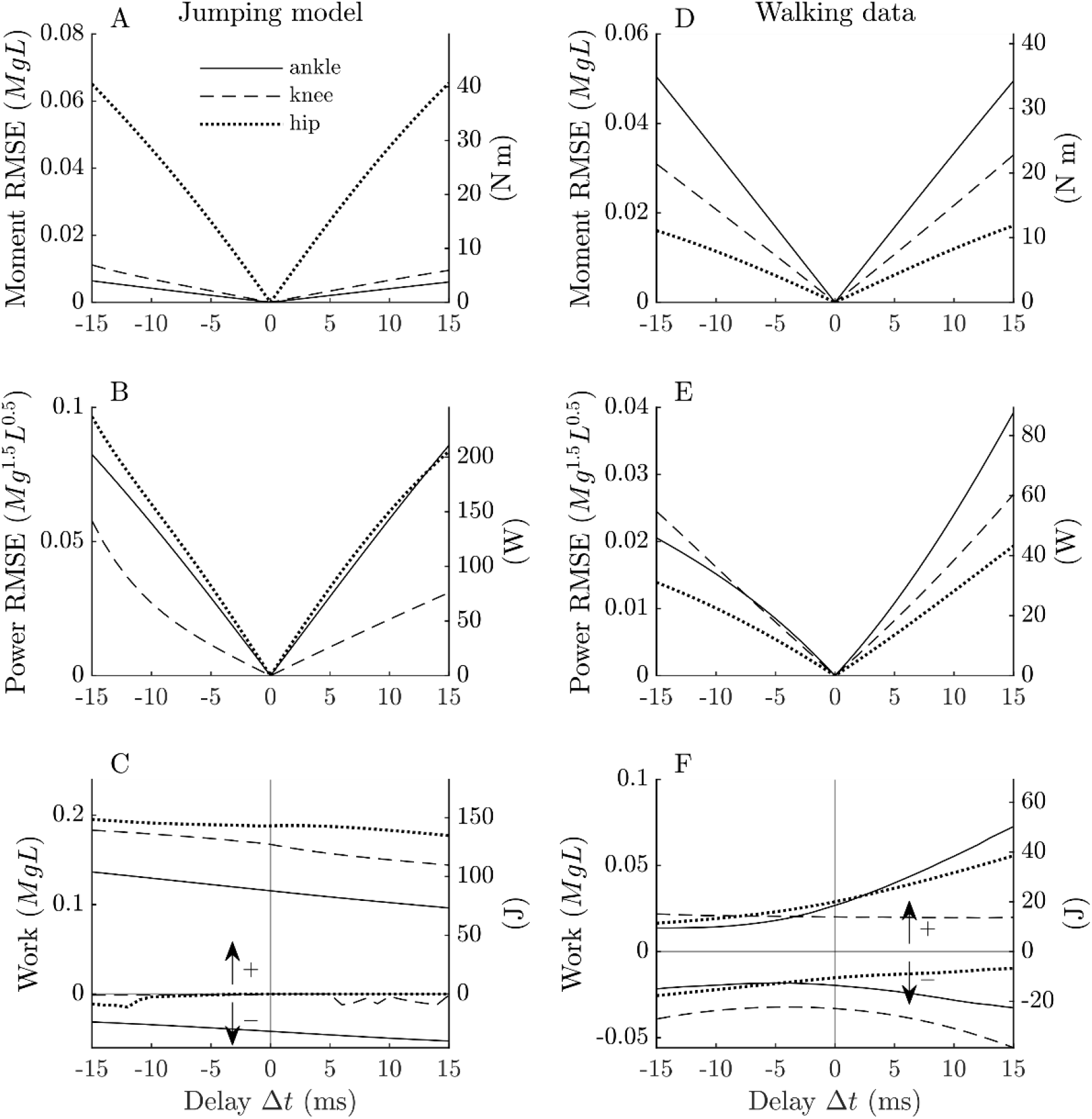
Summary of inverse dynamics errors due to mis-synchronization *Δt* for simulation of squat jumping (left column) and experimental data of human walking (right column), for ankle (solid), knee (dashed) and hip (dotted) joints, where mis-synchronization *Δt* is defined positive for forces lagging motion. Errors are shown as root-mean-square error (RMSE) for joint moments (Moment RSME, first row) and joint powers (Power RMSE, second row. Also shown is positive and negative mechanical work estimated for each joint (Work, third row), calculated as the time integral of the regions of positive and negative power over the take-off phase (jumping model) and over one complete stride (walking data). Here *Δt* = 0 represents the nominal reference case, for comparing how work is sensitive to non-zero mis-synchronization *Δt*. Jumping simulation data is illustrated in Fig. 1, walking data in Fig. 2. Left-hand vertical axes are in normalized units and right-hand axes are SI-units.

### Jumping simulation

For the jumping simulation, (Fig., 3 left column) the sensitivities of the joint moment RMSE to small delays around zero, for the ankle, knee and hip joints were 0.31, 0.52 and 3.78 N-m/ms, respectively (1.3, 2.2 and 16.0 non-dimensionalized units, from linear regression slopes). In other words, each millisecond of mis-synchronization resulted in an incremental error of 3.8 N-m in hip joint moment. The sensitivities for ankle, knee and hip joint power RMSE were 14.5, 5.3 and 16.7 W/ms, respectively (19.0, 7.0 and 22.0 non-dimensional). Especially for the hip joint, power as a function of time varied greatly as a function of delay (Fig. 1). In terms of positive and negative work done during the take-off phase (Fig. 3, left bottom), mis-synchronization also caused substantial differences compared to the nominal case, in particular for the ankle joint. For negative delays, net mechanical work was overestimated and for positive delays it was underestimated. For a +5 ms mis-synchronization in the jumping simulation data, the positive ankle work (defined as the time integral over the intervals where ankle power was positive) was underestimated by about 6%, at 83 J compared to 88 J in the nominal case.

### Experimental walking data

The errors due to mis-synchronization were also substantial for experimental walking data (Fig. 3, right column). The shape of moment and power curves differed substantially between positive and negative delays. Both joint moments and powers were generally smaller for negative delay than for positive delay (Fig. 2). Similar to the simulation results, the root-mean-square error in both joint moment and power increased approximately linearly with delay magnitude (Fig. 3, right column, top and middle). Around zero delay, the sensitivities of the joint moment RMSE for the ankle, knee and hip joints were 2.3, 1.5 and 0.8 N-m/ms, respectively (10.9, 6.9 and 4.0 non-dimensional). The sensitivities for ankle, knee and hip joint power RMSE were 4.4, 3.6 and 2.6 W/ms, respectively (6.4, 5.2 and 3.8 non-dimensional). For example, a +5 ms mis-synchronization in walking data yielded an RMSE of about 12 N-m in ankle joint moment and 27 W in ankle power, corresponding to respectively 29% and 58% of the total root-mean-squared signal at zero delay.

### Residual force and moment

As a separate measure of error, we quantified the residual force and moment for each delay, for both the jumping simulation and the walking data (Fig. 4). The residuals result from imperfect data being propagated sequentially through dynamical equilibrium equations for each body segment, typically ending with a non-zero force and moment acting from the sky on the top-most body segment. The residual force equals the difference between the sum of the forces acting on the model (ground reaction force and gravity) and the rate of change of total linear momentum of the model (based on rigid segment kinematics). The residual moment was computed about the whole-body center of mass (COM), as the difference between the sum of torques about the COM (based on ground reaction force and moment) and the rate of change of angular momentum about the COM (based on segment kinematics), where the location of the COM was based on segment kinematics. (The residual moment about any other point, such as COM of trunk, can be found by adding the moment of the residual force about that point.) Both the root-mean-square error (with respect to the ideal case of zero) of the residual force and that of the residual moment increased approximately linearly with mis-synchronization magnitude, for both the jumping simulation and the walking data (Fig. 4). For the jumping simulation, the sensitivities of the RMSE value of the residual force and moment around zero delay were 18.6 N/ms and 4.0 N-m/ms, respectively (74.2 and 17.1 non-dimensional). Consequently, for even a modest delay of 5 ms, the RMSE residual force was 11% of body weight, an error far greater than a typical force plate’s precision.

**Fig. 4.**
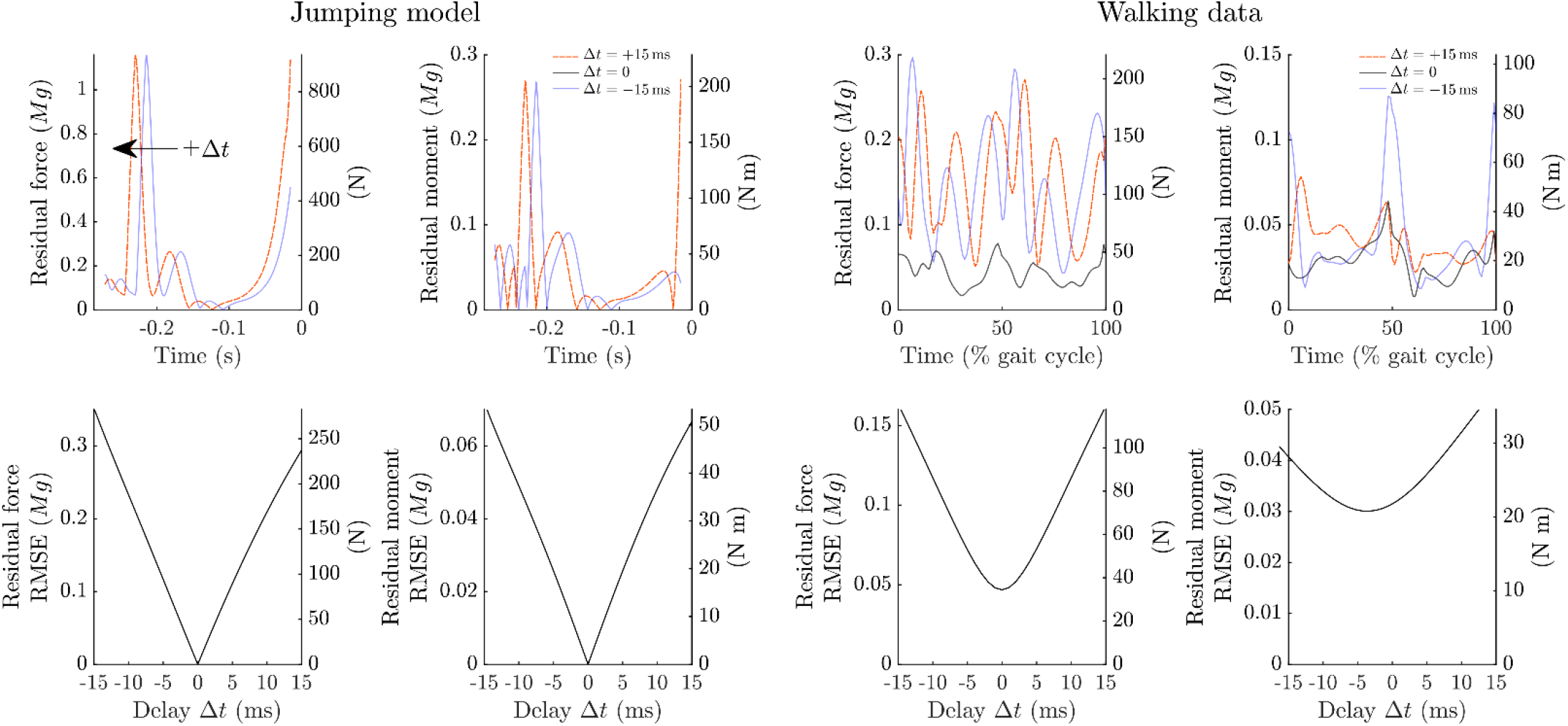
Residual forces and moments from (left) jumping simulation and (right) human walking data. Shown are (top row) residual forces and moments vs. time, for *Δt* = 0 and ±15 ms (positive delay, red dashed; negative delay, blue). Mis-synchronisation delay *Δt* is defined as positive for forces lagging motions. For the jumping simulation, *Δt* = 0 equals perfect synchronization, yielding zero residual forces and moments. For empirical walking data, *Δt* = 0 corresponds to a nominal synchronization (obtained by correcting an 8 ms lag in our force data), but still has non-zero residual forces due to other imperfections in the measurements and model. Left-hand vertical axes are in normalized units and right-hand axes are in SI-units.

For the walking data, the residual force and moment were non-zero at zero delay, because of modeling error and other errors present in the experimental data. In the linear portion of the curve, the increase in RMS of the residual force and moment per unit time was 6.1 N/ms and 0.8 N-m/ms, respectively (26.8 and 3.8 non-dimensionalized). This error sensitivity was almost 1% of body weight per ms of mis-synchronization, increasing to an error of 8% body weight at +5 ms of delay. For the residual moment, the sensitivity was about 1% of peak ankle torque per ms of mis-synchronization. Note that the residual force and moment had a minimum close to 0 ms delay, suggesting that the latency within our experimental set-up was corrected reasonably well.

### Effect of walking speed

We repeated the error analyses for a range of walking speeds (Fig. 5), to explore the error introduced by a mis-synchronization of +5 ms. Based on our theoretical analysis, errors should increase with movement amplitude and frequency, both of which increase with walking speed. We found that root-mean-square error between the nominal and the mis-synchronized case increased dramatically with walking speed for both joint moments and powers (Fig. 5A & B) and for residual forces and moments (Fig. 5C & D). For example, the error due to 5 ms of mis-synchronization in the ankle moment RMSE, increased from 6.6 N-m at a walking speed of 1.1 m/s to 16.8 N-m at 2.2 m/s, about a 250% increase in error. Over the same range of speeds, ankle power RSME increased even more dramatically from 9.0 W to 51.9 W, a greater than five-fold change. A less stark but still large increase was observed in the residual force RMSE, up to 11% body weight at 2.2 m/s, whereas residual moment RMSE was less greatly affected. Overall, errors due to mis-synchronization appear to be greatly exacerbated at faster movements, as expected from the theoretical analysis.

**Figure 5.**
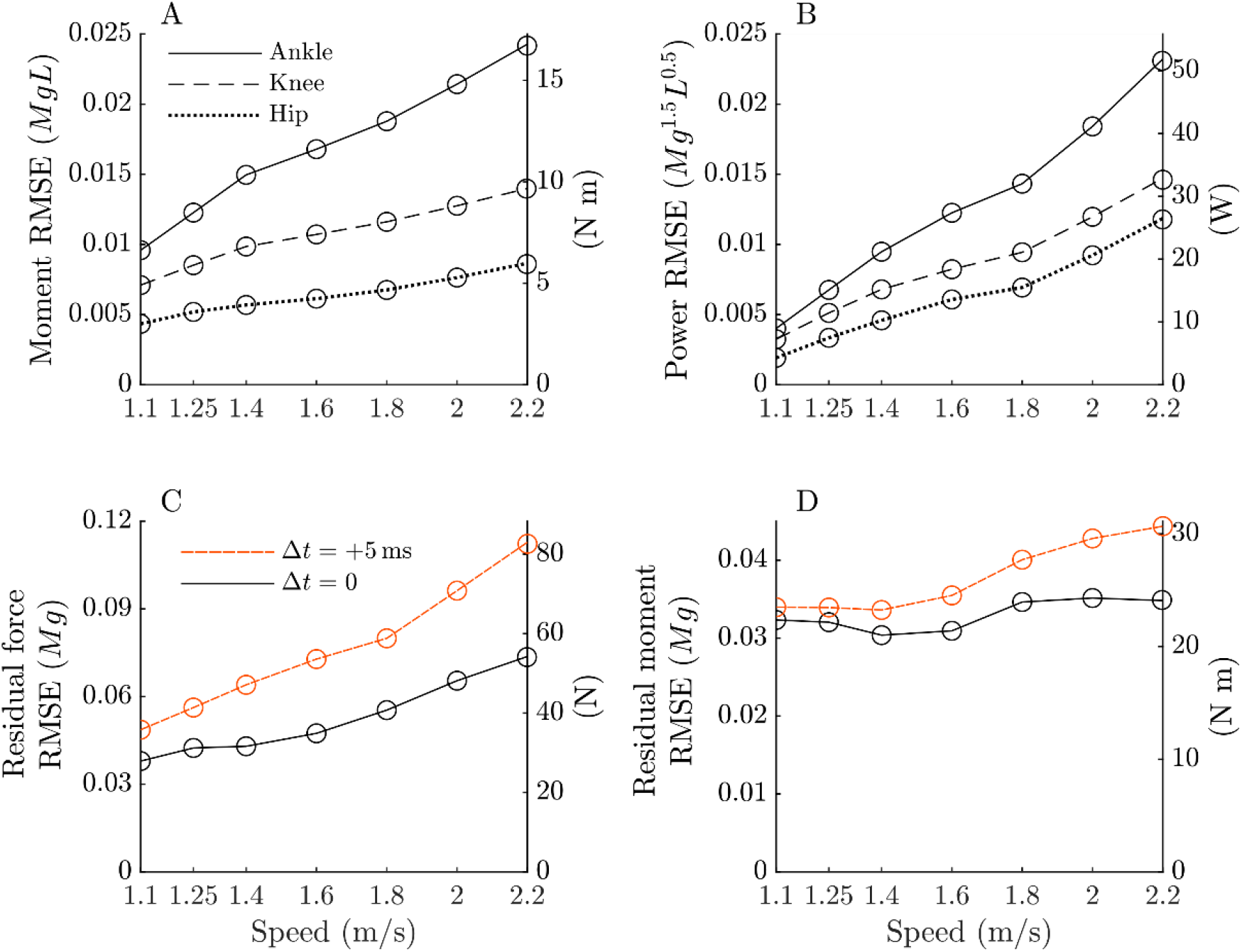
Inverse dynamics errors and residual forces and moments vs. walking speed, for data mis-synchronized by *Δt* = +5 ms about nominal. (A) Joint moment root-mean-square errors (RMSE) vs. walking speed, (B) Joint power RMSE errors vs. walking speed, for ankle, knee, and hip. (C) Residual force RMSE and (D) Residual moment RMSE vs. walking speed, for *Δt* = +5 ms (red, dashed) and *Δt* = 0 (black, solid) ms. Mis-synchronization delay was defined positive for force lagging motion data, where *Δt* = 0 corresponds to nominal synchronization. RMSE of (A) joint moments and (B) powers were quantified from the difference in results for *Δt* = +5 ms with respect to *Δt* = 0 (see Fig. 3). RMSE of (C) residual forces and (D) residual moments were quantified from the difference to the ideal case of zero residuals (see Fig. 4). Left-hand vertical axes are in normalized and right-hand axes are in SI-units.

## Discussion

We quantified the effect of mis-synchronization between force and motion capture data, on the results of inverse dynamics analysis. We did this in three ways, using a conceptual model, data from a human-like simulation of jumping, and experimental data of normal human walking. In all three cases, we found that mis-synchronization resulted in greater root-mean-square errors for both joint moments and powers. Here we discuss the types and the magnitudes of the errors that can occur with mis-synchronization, and provide some guidelines for anticipating such errors in general movements.

For both jumping and walking, the root-mean-square errors increased approximately linearly with mis-synchronization delay, as quantified by standard inverse dynamics outputs (Fig. 3) and by residual forces (Fig. 4). The two tasks differed in terms of where and when the largest errors occurred. In the jumping simulation, joint moment RMS errors were largest at the hip joint and smallest at the ankle joint. At +5 ms of mis-synchronization, the ankle joint moment RMSE was about 5% of peak ankle torque, whereas the hip joint moment RMSE was about 10% of peak hip torque. Conversely, joint power RMSE was similar across joints (Fig. 3). For the walking data, the errors were largest at the ankle joint and smallest at the hip joint. At +5 ms of mis-synchronization, ankle power RMSE was 27 W (58% of total ankle power RMS of the nominal data) whereas hip power RMSE was 14 W (42% of total hip power RMS of the nominal data). When averaged across joints, the moment RMS errors were of similar magnitude between the tasks, whereas the RMS power errors were about twice as large for the jumping simulation (Fig. 3). This finding is consistent with the analysis showing that errors in power are especially sensitive to higher frequency content of the signal, as jumping is a more explosive movement than walking. In terms of timing, there were no clear patterns in the error signal. It appeared that the timing of the greatest errors in moments and powers (Fig.1 and Fig. 2), and even the sign of the errors, simply depend on the specifics of the task. However, despite these complications, the overall root-mean-square errors (Fig. 3) did vary approximately linearly with mis-synchronization, as predicted analytically.

The analysis provides some insight into the effect of mis-synchronization on locomotion studies. The theoretical analysis shows that errors should increase sharply with motions of high frequency and high amplitude (specifically fourth power of frequency and square of amplitude, in Eq. 2). This leads to the expectation of greater adverse effects of mis-synchronization at higher walking speeds, which was indeed observed in the experimental data. Over the range of walking speeds investigated here (1.1 – 2.2 m/s), the RMS errors in ankle joint moment at +5 ms delay more than doubled across the range of speeds, whereas the RMS difference in ankle joint power increased by more than a factor of five. In addition, residual force almost tripled, up to 11% body weight at 2.2 m/s. At that speed, a mis-synchronization of +5 ms caused RMS errors in ankle moment and power of 34% and 86%, respectively. These errors are of a similar order as those previously reported for running (Günther et al., 2005), where a 5 ms mis-synchronisation yielded a 100% error in the work done by the ankle and hip joints during the contact phase at a speed of 4.8 m/s.

Similar error trends would be expected for movements other than those studied here. The actual errors depend on the specifics of the task, but as before, force errors should increase linearly with the mass and motion amplitude of the body segment, and with the third power of the motion frequency (Eq. 1). Joint power errors should increase still faster, with the square of the motion amplitude and the fourth power of frequency (Eq. 2). We therefore expect relatively slow and small motions such as standing balance to be relatively less sensitive to mis-synchronization, and more explosive motions to be quite sensitive.

The technique of introducing artificial mis-synchronization may be useful for evaluating the potential for errors due to mis-synchronization in any experimental dataset. The actual degree of mis-synchronization is usually unknown, but as applied by Günther (2005), artificial delays may be introduced to determine the effect on inverse dynamics results. This may be helpful for tasks other than locomotion. Where there is doubt about synchronization, the experimental force and motion data can be artificially shifted in time, and the inverse dynamics analysis repeated to determine the sensitivity.

An additional finding from our analysis was that residual forces are also sensitive to mis-synchronization, up to an error of 11% body weight at the highest walking speed. Residuals are due to erroneous inputs to the dynamical equilibrium equations, and in the standard iterative inverse dynamics procedure appear as non-zero residual forces and moments apparently acting from the sky on the top-most body segment (Faber et al., 2018). Non-zero residuals are generally observed in any experiment, and arise from imperfect data and imperfect modeling, including errors in force, motion, and body anthropometry. We found that mis-synchronization also contributes to residual forces (Fig. 4). In fact, as expected theoretically (Eq. 1), the residual magnitudes increased approximately linearly with mis-synchronization. In the present analysis of experimental data, we also found the magnitudes of residuals to be smallest near putative “zero delay.” If the other sources of experimental error are small, it is possible that residual forces can be helpful for identifying the presence or sensitivity of mis-synchronization.

We next suggest how motion-force synchronization should be performed. Commercial biomechanics laboratory equipment usually provides for precise synchronization of data capture, for example using motion cameras (e.g., Vicon motion systems ldt., UK) to trigger acquisition of force plate data. This does not necessarily account for processing latencies within the equipment, such as occurring within conventional force plates such as the 8 ms latency found in ours (Bertec Corp., Columbus, Ohio). We therefore find it helpful to synchronize data at the time of an applied physical stimulus, meaning when forces are exerted and are actually associated with motion. We believe that many force plates and most inertial measurement units introduce processing latencies, so that the outputted signals, even if recorded synchronously with motion capture, may still be delayed with respect to physical stimulus.

We have quantified the potential errors in inverse dynamics analysis due to mis-synchronization of motion and force data. Using a combination of theoretical analysis, computational simulation, and experimental testing, we found that quite substantial errors may result from rather small mis-synchronization. The errors are substantial for walking, and increase for faster or larger motions. It is often difficult to determine how much mis-synchronization is present in experimental data, but artificial shifting of the data, and then examining the resulting signals, including residual forces may give an indication. In addition, we find it critical to synchronize motion and force signals at the time of physical stimulus, and to compensate for processing latencies present in equipment. These tests and compensations can help ensure the accuracy of inverse dynamics analysis of experimental data.

## Acknowledgements

This work supported in part by the Natural Sciences and Engineering Research Council of Canada (NSERC Discovery and Canada Research Chair, Tier 1) and the Dr. Benno Nigg Research Chair in Biomechanics.

